# *Arabidopsis* LSH10 transcription factor interacts with the co-repressor histone deubiquitinase OTLD1 to recruit it to the target promoters

**DOI:** 10.1101/2022.07.30.502139

**Authors:** Mi Sa Vo Phan, Ido Keren, Phu Tri Tran, Moshe Lapidot, Vitaly Citovsky

**Affiliations:** Department of Biochemistry and Cell Biology, State University of New York, Stony Brook, NY 11794-5215; Department of Vegetable Research, Institute of Plant Sciences, Agricultural Research Organization, The Volcani Institute, 68 HaMaccabim Rd, P.O.B 15159, Rishon LeZion 7505101, Israel

**Author notes:** Current address (Ido Keren).

## Abstract

Histone ubiquitylation/deubiquitylation plays a major role in the epigenetic regulation of gene expression. In plants, OTLD1, a member of the ovarian tumor (OTU) deubiquitinase family, deubiquitylates monoubiquitylated histone 2B and represses the expression of genes involved in growth, cell expansion, and hormone signaling. Like many other histone-modifying enzymes, OTLD1 lacks the intrinsic ability to bind DNA. How OTLD1, as well as most other known plant histone deubiquitinases, is recruited specifically to the promoters of its target genes remains unknown. Here, we show that *Arabidopsis* transcription factor LSH10, a member of the ALOG protein family, interacts with OTLD1 in living plant cells. Loss-of-function LSH10 mutations relieve the OTLD1-promoted transcriptional repression of the target genes, resulting in their elevated expression, whereas recovery of the LSH10 function results in down-regulated transcription of the same genes. We then show that LSH10 associates directly with the target gene chromatin as well as with the specific DNA sequence motifs in the promoter regions of the target genes. Furthermore, in the absence of LSH10, the degree of H2B monoubiquitylation in the target promoter chromatin increases. Hence, our data suggest that OTLD1-LSH10 acts as a co-repressor complex, in which LSH1 recruits OTLD1 to the target gene promoters, potentially representing a general mechanism for recruitment of plant histone deubiquitinases to the target chromatin.

## Introduction

Growing in a dynamic environment, plants must deal with numerous biotic and abiotic stress sources. Histone modifications that underly epigenetic regulation of gene expression help plants to adapt to environmental changes and serve as an environmental memory of the transcription (1). Histone modifications not only regulate gene expression in response to diverse environmental signals such as stress (2), pathogen attack (3), temperature (4, 5), and light (6) but play an important role in plant development and morphogenesis [reviewed in (4, 7-10)].

Covalent modifications of histones can activate or repress transcription by altering the chromatin structure (11). However, because histone-modifying enzymes often do not possess a DNA binding ability, they are not sufficient to trigger transcriptional regulation of the specific target genes. To achieve this regulation, histone-modifying enzymes are thought to function in complexes with transcription factors that contain DNA-binding domains and, thus, can provide the DNA binding capacity to the histone modifier-transcription factor complex and thereby recruit histone-modifying enzymes to the target promoters. Indeed, in plants, different transcription factors have been shown to recruit such diverse histone modifiers as histone methyltransferases, histone acetyltransferases, histone demethylases, and Polycomb repressive complexes that promote histone trimethylation and monoubiquitylation. Yet, whether transcription factors can also mediate the recruitment of plant histone deubiquitinases, one of the major types of histone-modifying enzymes, remains unknown.

OTLD1, which belongs to the ovarian tumor (OTU) deubiquitinase family, represents one of the few characterized plant histone deubiquitinases (12, 13). Our previous studies have shown that OTLD1 mainly functions as a transcriptional co-repressor (although, at least in one case, OTLD1 facilitated transcriptional activation of its target gene) by associating with the target chromatin and deubiquitylating histone 2B (H2B) at the occupied regions, thereby promoting the erasing or writing of euchromatic histone acetylation and methylation marks (14, 15). Consistent with this notion, OTLD1 can interact and crosstalk with the histone demethylase KDM1C to coordinate histone modification and transcriptional regulation of the target genes (16, 17). But how is OTLD1 recruited to the target gene promoters? To address this question, we endeavored to identify a putative DNA-binding protein that recognizes OTLD1 and targets it to the sites of its biological function.

Here, we report that LSH10, a member of the ALOG (Arabidopsis LSH1 and Oryza G1) protein family, interacts with OTLD1 and participates in transcriptional repression of the OTLD1 target genes *OSR2, WUS, ABI5*, and *ARL*. The expression of these genes was elevated in the *LSH10* loss-of-function mutants and suppressed in the gain-of-function lines. Specific interaction between LSH10 and OTLD1 was demonstrated by two independent fluorescence imaging techniques in living plant cells. The binding of LSH10 to the chromatin and DNA sequences of the *OSR2, WUS, ABI5*, and *ARL* promoters was also demonstrated. The *LSH10* loss-of-function plants also exhibited enhanced H2B monoubiquitylation in the target promoter chromatin. Taken together, these observations suggest that LSH10 functions as a transcription factor that interacts with the OTLD1 co-repressor to recruit it to the promoters of the target genes.

## Results

### LSH10 is a nuclear protein that interacts with OTLD1

To gain better insight into possible mechanisms by which plant histone deubiquitinases are recruited to their target chromatin, we searched for proteins that interact with OTLD1. A truncated OTLD1 was used as bait to screen the Arabidopsis yeast two-hybrid protein interaction library (18), and the LSH10 protein was identified as a putative interaction partner of OTLD1 (not shown). We then examined the subcellular localization of LSH10, which was tagged with CFP and transiently expressed in *N. benthamiana* leaf tissues together with a free YFP reporter that partitions between the cell cytoplasm and the nucleus, conveniently visualizing and identifying both subcellular compartments. Fig. 1A shows that LSH10-CFP accumulated in the cell nucleus of the cells whereas, as expected, the free YFP fluorescence was nucleocytoplasmic. This nuclear localization of LSH10 is consistent with its interaction with OTLD1, a histone deubiquitinase that functions in the cell nucleus (16).

**Fig. 1.**
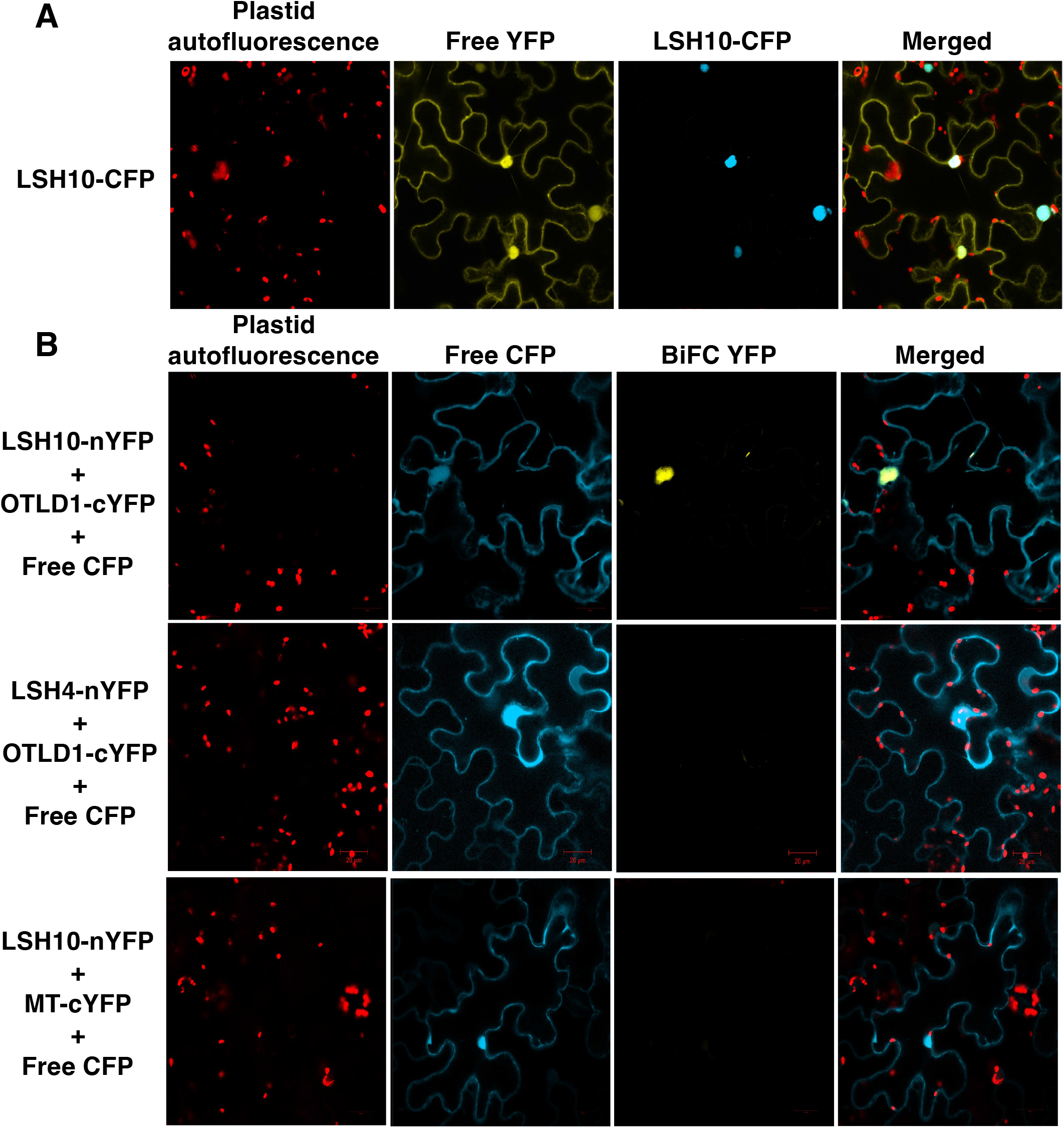
LSH10 subcellular localization, and specific interaction of LSH10 with OTLD1 *in planta* detected by BiFC. (**A**) Nuclear localization of LSH10. The LSH10 protein tagged with CFP (LSH10-CFP) and free YFP were transiently co-expressed in *N. benthamiana* leaves. (**B**) The BiFC assay of the interaction between LSH10 and OTLD1. The indicated combinations of proteins tagged with nYFP and cYFP were transiently co-expressed in *N. benthamiana* leaves. CFP signal is in cyan; YFP signal is in yellow; merged CFP and YFP signals are in cyan-yellow, and chlorophyll autofluorescence is in red. All images are single confocal sections.

Next, the physical interaction of LSH10 with OTLD1 was studied in planta using two independent approaches, bimolecular fluorescence complementation (BiFC) and fluorescence resonance energy transfer (FRET). BiFC experiments shown in Fig. 1B detected a strong fluorescent signal of the reconstituted YFP in the cells co-expressing both LSH10 and OTLD1, indicating protein interaction. This interaction was specific as no BiFC signal was observed in the cells co-expressing either OTLD1 and LSH4, a homolog of LSH10, or LSH10 and an unrelated plant viral protein MT (Fig. 1B). Furthermore, the interacting proteins colocalized with the nuclear portion of the coexpressed free YFP reporter, indicating that the OTLD1-LSH10 complexes were located in the cell nucleus, the expected subcellular site of their function (Fig. 1B). Our FRET experiments—using LSH10 tagged with GFP as donor fluorophore and OTLD1 tagged with RFP as acceptor fluorophore—confirmed and extended the BiFC findings. We used two variations of the FRET method, sensitized emission (SE-FRET) and acceptor bleaching (AB-FRET) (19). In SE-FRET, protein interaction results in the transfer of the excited state energy from the GFP donor to the RFP acceptor without emitting a photon, producing the fluorescent signal with an emission spectrum similar to that of the acceptor. AB-FRET, on the other hand, detects and quantifies protein interaction from increased emission of the GFP donor when the RFP acceptor is irreversibly inactivated by photobleaching. Fig. 2A summarizes the results of the SE-FRET experiments, in which the cell nuclei were simultaneously recorded in all three, i.e., donor GFP, acceptor RFP, and SE-FRET, channels and used to generate images of SE-FRET efficiency illustrated in a rainbow pseudo-color. This color scale, i.e., transition from blue to red, indicates an increase in FRET efficiency from 0 to 100%, which corresponds to the degree of protein-protein proximity during the interaction. The SE-FRET signal observed in the cell nuclei following coexpression of LSH10 and OTLD1 was comparable to that generated in positive control experiments which expressed the translational acceptor-donor RFP-GFP fusion. Negative controls, i.e., coexpression of OTLD1-RFP with LSH4-GFP or free RFP with LSH10-GFP, produced no SE-FRET signal (Fig. 2A). The FRET data were quantified using AB-FRET (Fig. 2B, C) by recording the cell nuclei in the donor GFP channel before and after RFP photobleaching and displayed in pseudo-color to visualize the change in GFP fluorescence. Fig. 2B shows that photobleaching of the RFP acceptor completely blocked its fluorescence in all protein coexpression combinations tested. Following this photobleaching, two protein combinations showed an increase in the GFP donor signal, i.e., LSH10-GFP coexpressed with OTLD1-RFP and the RFP-GFP fusion positive control. In contrast, the negative controls, i.e., LSH4-GFP coexpressed with OTLD1-RFP and LSH10-GFP coexpressed with free RFP, elicited no increase in the GFP fluorescence (Fig. 2B). Quantification of these data demonstrated that the donor dequenching (%AB-FRET) of 13% observed following LSH10-OTLD1 coexpression was statistically significant and overall comparable to the maximal %AB-FRET of 30% achieved with RFP-GFP. Both negative controls displayed 0% dequenching (Fig. 2C). Collectively, the data in Fig. 2 indicate that LSH10 interacts with OTLD1 within living plant cells, that the interacting proteins accumulate in the cell nucleus, and that, in the LSH10-OTLD1 complex, the proteins are within <10 nm from each other, the effective range of protein interactions detected by FRET (20).

**Fig. 2.**
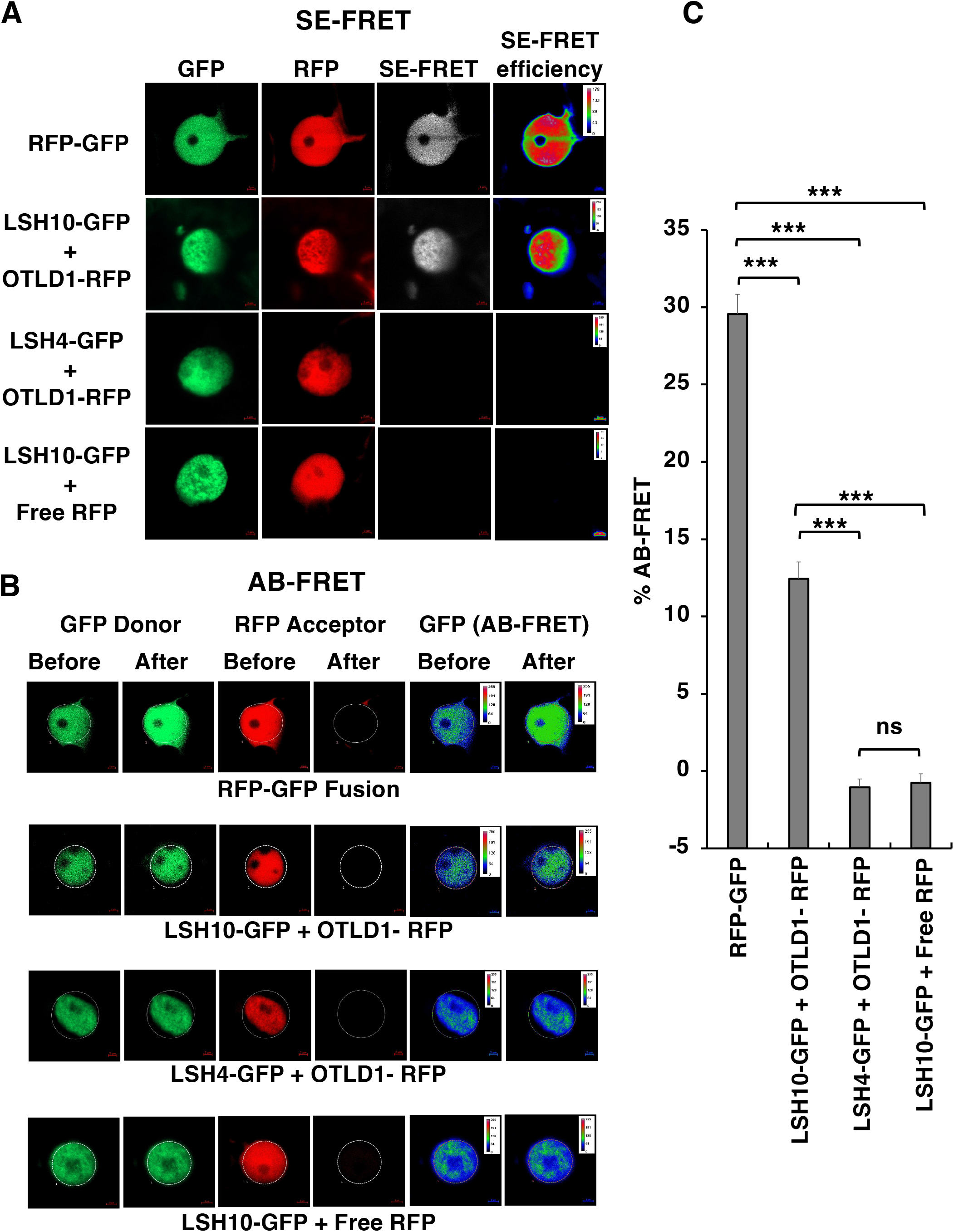
Specific interaction of LSH10 with OTLD1 *in planta* detected by FRET. The indicated combinations of proteins tagged with GFP (energy donor) and RFP (energy acceptor) were transiently co-expressed in *N. benthamiana* leaves. (**A**) SE-FRET. Images were collected from the three detection channels, i.e., donor (GFP), acceptor (RFP), and raw SE-FRET. The SE-FRET efficiency images represent the calculated corrected image after subtraction of spectral bleed-through and are presented in a rainbow pseudo-color scale, in which red denotes the highest SE-FRET signal and blue denotes the lowest signal. (**B**) AB-FRET. Images were collected from the two detection channels, i.e., donor (GFP) and acceptor (RFP), before and after photobleaching. The circle delineates a region that was photobleached and its fluorescence measured. The AB-FRET is presented in a rainbow pseudo-color scale, in which red denotes the highest GFP signal and blue denotes the lowest signal. (**C**) Quantification of AB-FRET. Data measured in panel B were used to calculate %AB-FRET. Error bars represent the mean for N = 13 cells for each measurement. Numerical values of individual data points are listed in Table S2. Differences between mean values assessed by the two-tailed t-test are statistically significant for the p-values * p < 0.05, ** p < 0.01, and *** p < 0.001; P ≥ 0.05 are not statistically significant (ns).

### LSH10 has structural features of a transcription factor

LSH10 is a 177-amino acid residue protein (Fig. 3A) encoded by the Arabidopsis At2G42610 gene. It belongs to a 10-member family of Arabidopsis ALOG (Arabidopsis LSH1 and Oryza G1) protein family (Fig. S1), many of which remain uncharacterized. The members of this family carry a highly conserved ALOG domain (also known as DOMAIN OF UNKNOWN FUNCTION 640 / DUF640) located in the center of the protein molecule. This domain is composed of 4 all-α helices, a zinc ribbon insert structure, and a nuclear localization signal (NLS) (Fig. 3). ALOG is predicted to act as a DNA binding domain and belongs to the tyrosine recombinase/phage integrase N-terminal DBD superfamily (21), in which the ALOG domain members, unlike the tyrosine recombinase members, contain a conserved zinc ribbon insert located between helices 2 and 3 with highly conserved positively charged residues at its N-terminus and the “HxxxC’’ and “CxC” motifs. This region can provide additional molecular contacts unique to the ALOG domain to participate in binding to DNA. The conserved ALOG sequences (Fig. 3) and our prediction of the DNA-binding amino acid residues using three methods, DRNApred (22), DP-Bind (23), and DISPLAR (24), indicate that LSH10 may associate with DNA via hydrogen bonding and ionic interactions with multiple conserved solvent-accessible basic residues (25) in helix-1, helix-3, helix-4, and in the zinc ribbon or with a conserved acidic residue (25) in helix-1, whereas the conserved hydrophobic residues in all four helices likely stabilize the core tetra-helical fold (26) in the target DNA molecule (Fig. 3). Thus, the sequence analysis of LSH10 indicates that this protein binds DNA, consistent with its proposed activity as a transcription factor.

**Fig. 3.**
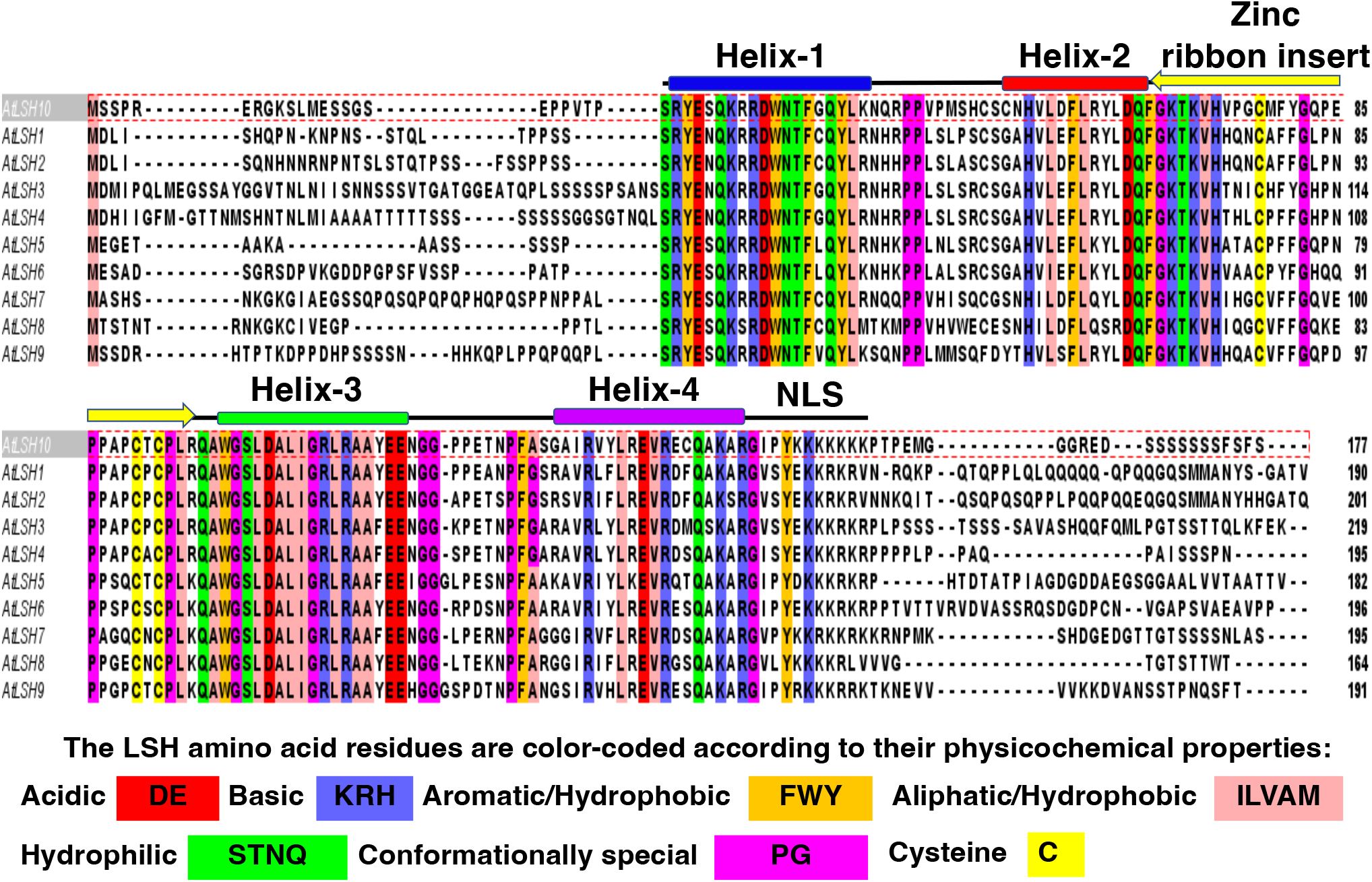
Structure of LSH10 (ALOG domain). Sequence alignment of the nine Arabidopsis LSH (AtLSH) protein family members and conserved features of their ALOG domain. Sequences are color-coded based on their conservation at 100% consensus. The coloring scheme and the conserved amino acids are indicated below the alignment. The highly conserved ALOG domain includes a zinc ribbon insert structure, four helices, and an NLS.

### LSH10 is a transcriptional repressor of the OTLD1 target genes

The interaction of LSH10 with OTLD1 and its potential DNA binding ability suggest that LSH10 may function as a transcription factor that recruits the OTLD1 co-repressor to its target genes. In this scenario, LSH10 should function in complex with OTLD1 and, thus, repress at least a subset of the target genes repressed by OTLD1. Our previous study indicated that OTLD1 is involved in the transcriptional repression of five genes *OSR2, WUS, ABI5, ARL*, and *GA20OX* (15). Thus, we examined the effect of *LSH10* loss-of-function mutations on the transcription of these genes. To this end, two *Arabidopsis lsh10* T-DNA insertion lines (SALK_006965 and SK14678) were obtained from ABRC (www.arabidopsis.org/abrc/) and the homozygous mutant lines, designated *lsh10-1* and *lsh10-2*, were generated. The *lsh10-1* and *lsh10-2* mutants contained a single T-DNA insertion in the exon and 5’UTR of the *LSH10* gene, respectively (Fig. 4A). The RT-qPCR analysis showed that, in both *lsh10-1* and *lsh10-2* plants, the transcription of the *LSH10* gene was virtually abolished (Fig. 4B), confirming the loss of function of this gene in both mutant lines.

**Fig. 4.**
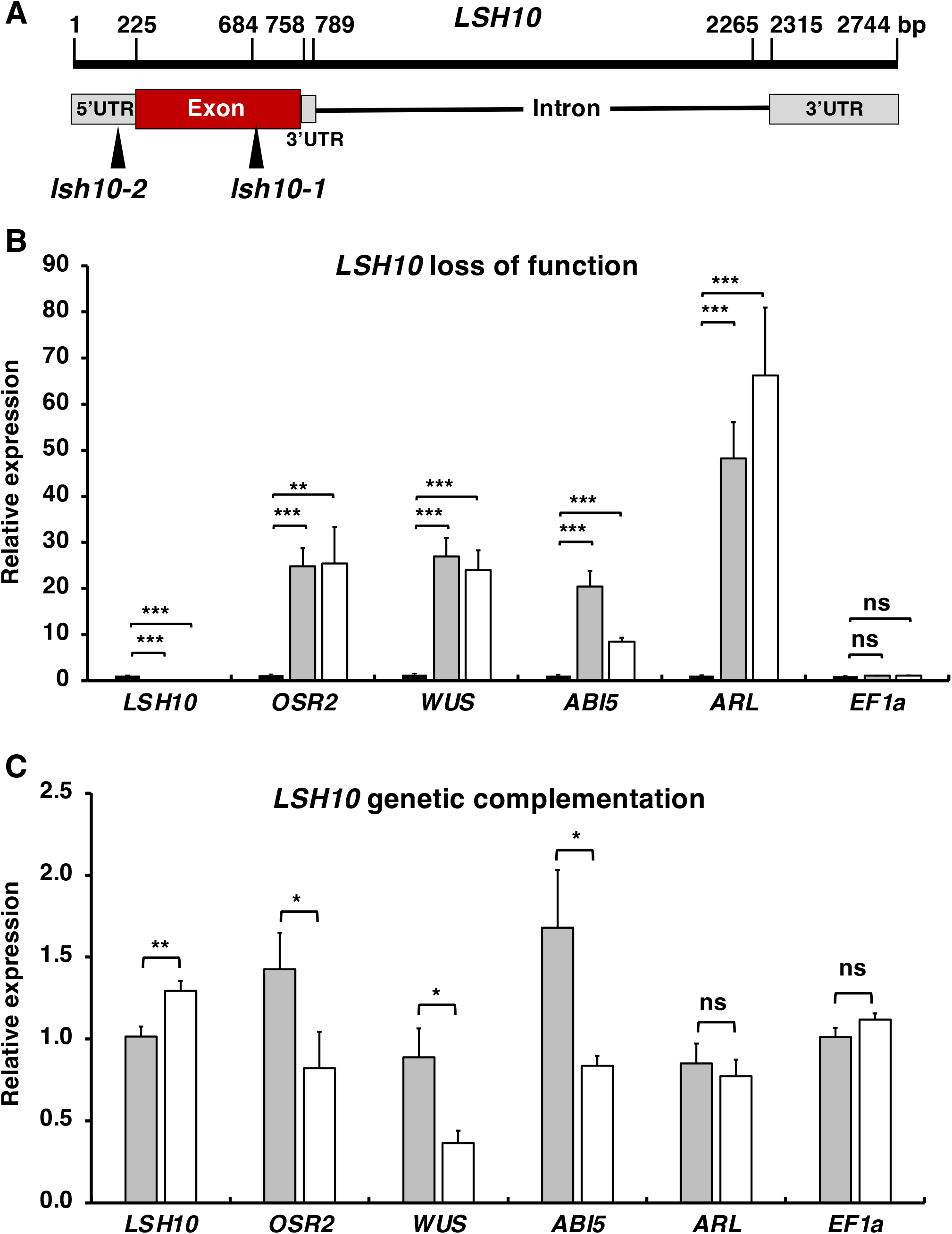
LSH10 is a transcriptional repressor of the OTLD1 target genes *OSR2, WUS, ABI5*, and *ARL*. (**A**) A schematic structure of the *LSH10* gene and its loss-of-function alleles *lsh10-1* and *lsh10-2*. The locations of the mutagenic T-DNA inserts in each of the alleles are indicated by arrowheads. (**B**) Increase in expression of the indicated target genes in the *lsh10-1*, and *lsh10-2* mutants. Wild-type plants, black bars; *lsh10-1*, gray bars; *lsh10-2*, white bars. (**C**) Transcriptional repression of the indicated target genes in genetically complemented *lsh10-1/LSH10-His6* plants. Wild-type plants, gray bars; *lsh10-1/LSH10-His6*, white bars. The increase in gene expression and transcriptional repression were analyzed by RT-qPCR with primers listed in Table S1. Error bars represent the mean for N = 8 biological replicates. Numerical values of individual data points are listed in Tables S3 and S4. Differences between mean values assessed by the two-tailed t-test are statistically significant for the p-values * p < 0.05, ** p < 0.01, and *** p < 0.001; P ≥ 0.05 are not statistically significant (ns).

Next, the amounts of transcripts of each of the *OSR2, WUS, ABI5, ARL*, and *GA20OX* genes were analyzed by RT-qPCR in the *lsh10-1* and *lsh10-2* plants and compared to the wild-type plants. Each of the *OSR2, WUS, ABI5*, and *ARL* genes displayed a substantial and statistically significant increase in expression in both loss-of-function lines (Fig. 4B). Specifically, the amounts of the *OSR2, WUS, ABI5*, and *ARL* transcripts were elevated ca. 20.8 to 21.4-fold, 21.08 to 18.79-fold, 18.39 to 7.61-fold and 43.85 to 60.2-fold in *lsh10-1* and *lsh10-2*, respectively. The expression of the internal reference gene *EF1a* was not significantly altered in any of the plant lines (Fig. 4B). Interestingly, we also observed a reduction in the expression of *GA20OX* (not shown), suggesting that regulation of this gene may involve other transcription factors.

Besides testing two different alleles of the *lsh10* loss-of-function mutant, we confirmed that derepression of the OTLD1 target genes resulted from the decrease in *lsh10* transcription by genetic complementation of one of the alleles, *lsh10-1*, with the wild-type *LSH10* coding sequence. We generated a transgenic *lsh10-1* line, *lsh10-1/LSH10-His6*, that expresses wild-type LSH10 protein tagged with hexahistidine. The resulting *lsh10-1/LSH10-His6* plants expressed the tagged *LSH10* at higher levels than the parental *lsh10-1* plants (compare Fig. 4C to Fig. 4B), these levels were comparable and even slightly, ca. 1.5-fold, higher than the levels of the endogenous *LSH10* transcript in the wild-type plants (Fig. 4C). In these genetically complemented plants (Fig. 4C), relative to the parental plants (Fig. 4B), we observed clear repression of all four target genes, i.e., *OSR2, WUS, ABI5*, and *ARL*, whereas no such repression was detected with the negative control *EF1a* gene (Fig. 4C). Noteworthy, the target gene repression in the *lsh10-1/LSH10-His6* plants was more pronounced not only relative to the parental, loss-of-function *lsh10-1* line (compare Fig. 4C to Fig. 4B) but, in most cases, also relative to the wild-type line with the native expression of the wild-type *LSH10* gene (Fig. 4C). Collectively, these observations indicate that LSH10 acts as a transcriptional repressor of most of the known OTLD1 target genes.

### LSH10 binds to the promoter DNA sequences and associates with the chromatin of the OTLD1/LSH10 target genes to deubiquitylate H2B

The proposed function of LSH10 as a transcriptional repressor that recruits the OTLD1 co-repressor to the target chromatin implies that LSH10 binds directly to the regulatory sequences of the gene regulated both by OTLD1 and LSH10. Thus, we examined whether LSH10 can bind the promoters of the *OSR2, WUS, ABI5*, and *ARL* genes directly, using the electrophoretic mobility shift assay (EMSA). We selected 2-4 conserved motifs of the intergenic regions of each of these genes (Fig. 5A, D) and used them as EMSA probes for interaction with a purified recombinant LSH10 tagged with GST (glutathione-S-transferase). Fig. 5B shows that each of these probes was recognized by LSH10 as detected by substantially reduced electrophoretic mobility of the GST-LSH10-probe complexes as compared to the free probe (lanes 3, 7, 11, 15, 19, 23, 27, 31, 35, 39 and lanes 1, 5, 9, 13, 17, 21, 25, 29, 33, 37 respectively). This binding was specific because it was not observed with GST alone (Fig. 5B, lanes 2, 6, 10, 14, 18, 22, 26, 30, 34, 38) and substantially reduced in the presence of competing amounts of unlabeled DNA, corresponding to each probe (Fig. 5B, lanes 4, 8, 12, 16, 20, 24, 28, 32, 36). Consistent with this binding specificity, not all selected motifs were recognized by LSH10 (e.g., Fig. 5B, lanes 40, 41, 42). Taken together, the EMSA experiments lend support to the idea that LSH10 functions as a DNA binding protein that recognizes sequence elements within the target gene promoters. That several diverse promoters are recognized by LSH10 suggests a “fuzzy” type of recognition that allows a single transcription factor to bind variable consensus DNA sequences (27).

**Fig. 5.**
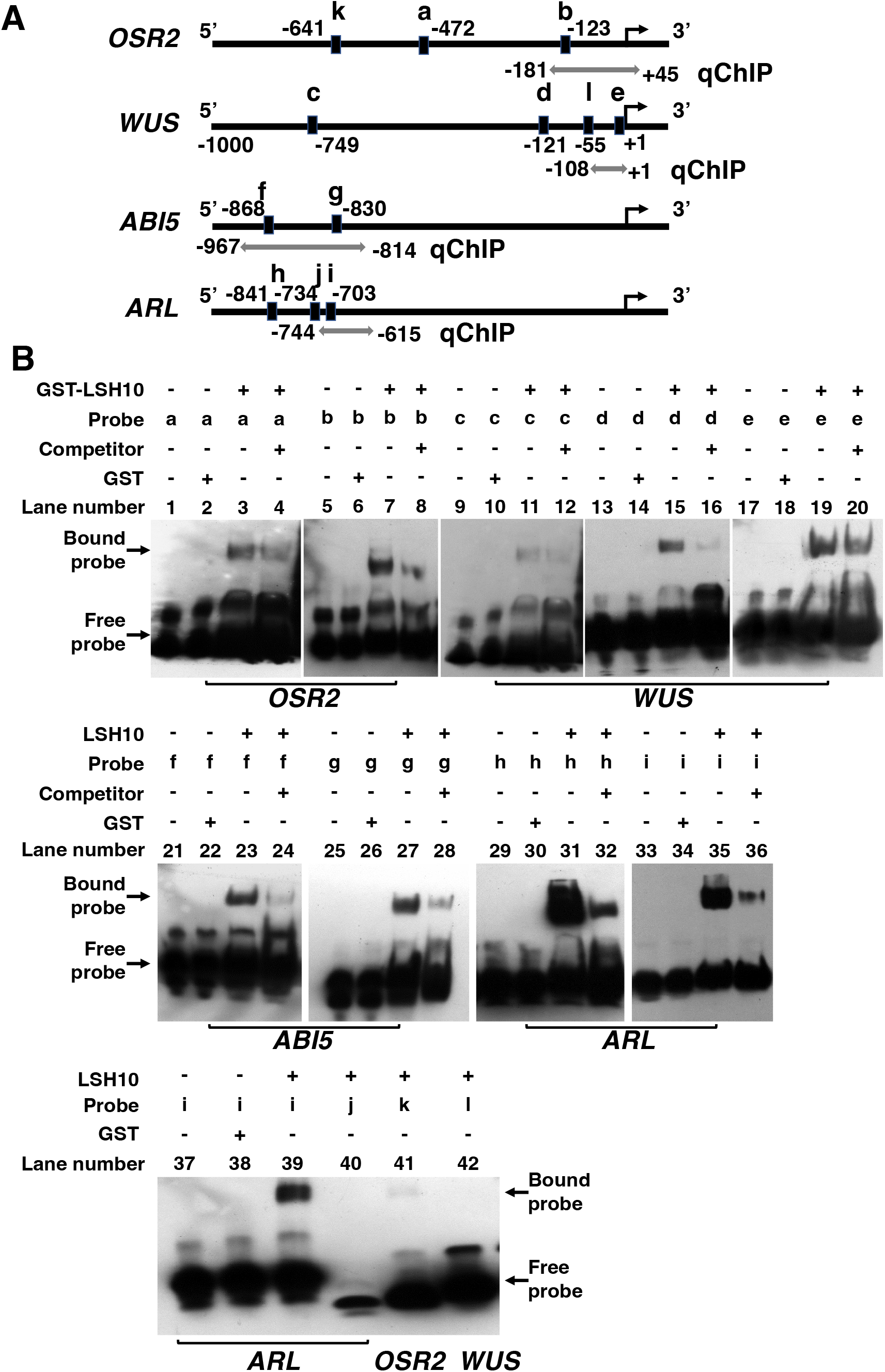
Binding of LSH10 to the promoter DNA sequences of the *OSR2, WUS, ABI5*, and *ARL* target genes. (**A**) A schematic of the *OSR2, WUS, ABI5*, and *ARL* gene promoters showing the locations of probe sequences for EMSA (denoted by black rectangles and letters “a”-”i” and detailed in Table S1) and primers for qCHIP (denoted by double-headed gray arrows and detailed in Table S1) relative to the translation initiation sites (bent arrows). (**B**) EMSA of LSH10 binding to promoter sequences of the indicated target genes. The composition of the binding mixtures is detailed above each lane of the gel image, with (+) or (-) signifying the presence or absence, respectively, of the indicated components. The electrophoretic mobilities of the LSH10-bound and free probes are indicated by arrows.

Next, we investigated the potential association of LSH10 with its target gene chromatin within plant cells, taking advantage of the fact that, in the *lsh10-1/LSH10-His6* line, LSH10 is tagged with an His6 epitope, allowing us to utilize quantitative chromatin immunoprecipitation (qChIP) to detect its presence. To correlate the physical association of LSH10 with the target chromatin and the binding of LSH10 to the target promoter sequence, the qChIP primers were designed to overlap several EMSA probes (Fig. 5A). Fig. 6A shows that, indeed, LSH10 was associated with the regions of the *OSR2, WUS, ABI5*, and *ARL* chromatin that contained the DNA sequences to which LSH10 was able to bind (Fig. 5). The amounts of immunoprecipitated LSH10-His6 were statistically significant yet varied between different target genes, potentially reflecting the different amounts of the LSH10/OTLD1-containing repressor complexes involved in the repression of each of these genes. Overall, these data demonstrate the ability of LSH10 to bind the conserved DNA motifs in the target gene promoters and to associate with the promoter chromatin of these genes.

**Fig. 6.**
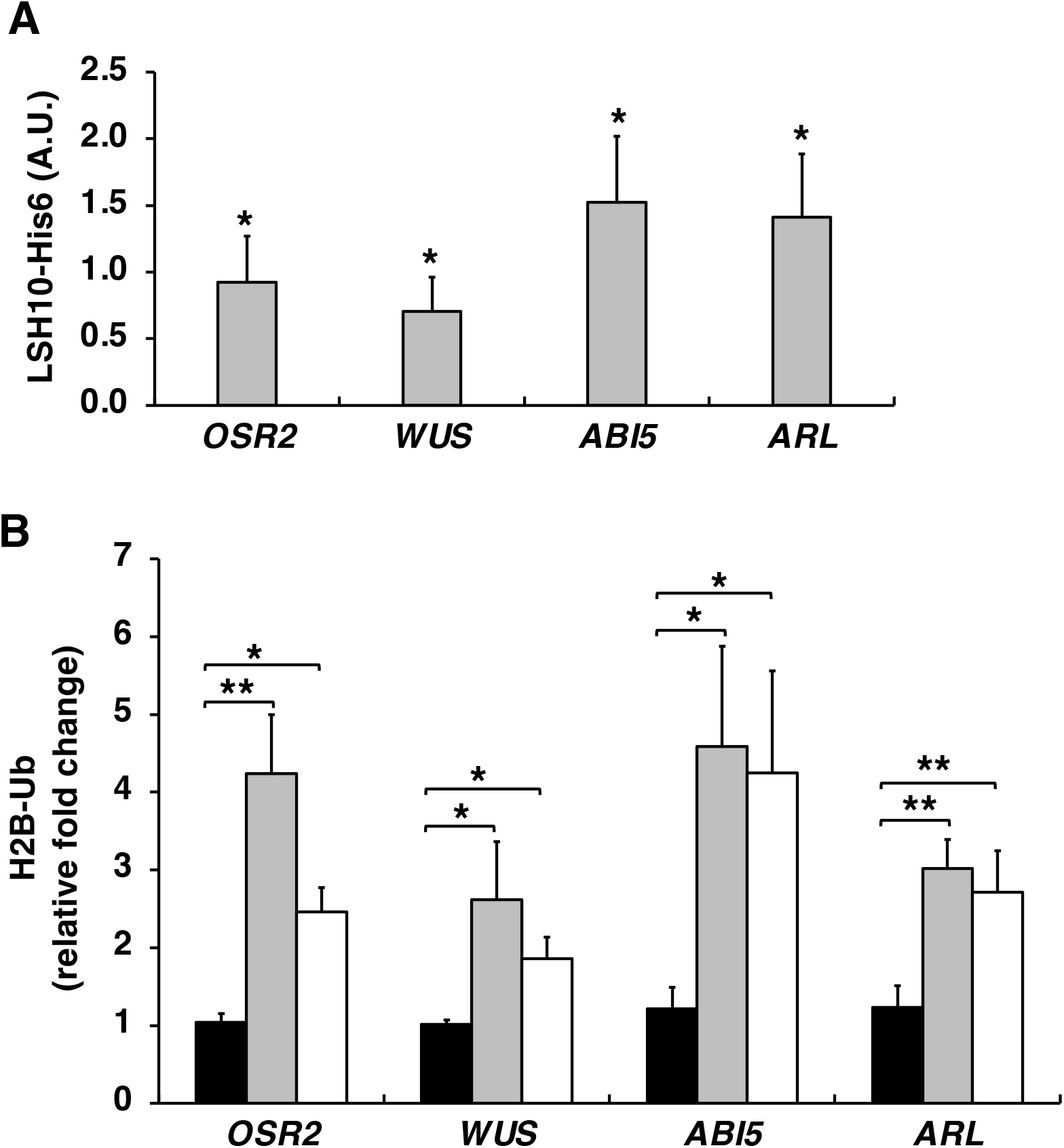
Association of LSH10 with and effects on ubiquitylation of the chromatin of the *OSR2, WUS, ABI5*, and *ARL* genes. (**A**) Association of LSH10-His6 with the chromatin of the indicated target genes in the *lsh10-1/LSH10-His6* plants. A.U., arbitrary units. (**B**) Increase in H2B monoubiquitylation of the *OSR2, WUS, ABI5*, and *ARL* promoter chromatin in the *lsh10-1* mutant plants. Wild-type plants, black bars; *lsh10-1*, gray bars; *lsh10-2*, white bars. Chromatin association of LSH10 and the degree of H2B monoubiquitylation were analyzed by qChIP with primers described in Fig. 5A and listed in Table S1. Error bars represent the mean for N>5 biological replicates with 3 technical repeats for each. Numerical values of individual data points are listed in Tables S5 and S6. Differences between mean values assessed by the two-tailed t-test are statistically significant for the p-values * p < 0.05 and ** p < 0.01.

Finally, we examined the notion that, if LSH10 recruits OTLD1 to deubiquitylate the target chromatin, such deubiquitylation will be reduced in the absence of LSH10. Thus, we used qChIP to analyze the promoter chromatin of the *OSR2, WUS, ABI5*, and *ARL* genes in the loss-of-function *lsh10-1* and *lsh10-2* mutants in and in the wild-type plants for the presence of monoubiquitylated H2B, known to be deubiquitylated by OTLD1 in these chromatin regions (16). Fig. 6B shows a significant degree of hyperubiquitylation of H2B in the *OSR2, WUS, ABI5*, and *ARL* chromatin in both mutants as compared to the wild-type plants. Specifically, the *OSR2* chromatin of *lsh10-1* and *lsh10-2* plants was monoubiquitylated on average 4.08-2.37-fold more than the wild-type *OSR2* chromatin, and monoubiquitylation of *WUS, ABI5* and *ARL* was increased 2.59-1.84-fold, 3.76-3.48-fold, and 2.46-2.20 -fold, respectively (Fig. 6B).

## Discussion

Epigenetic regulation of transcriptional outcomes heavily relies on the action of histone-modifying enzymes that function as writers and erasers of euchromatic and heterochromatic marks. Most of these enzymes, however, lack DNA binding capabilities and, therefore, are unable to recognize their target chromatin. Instead, this targeting most likely is mediated by DNA-binding transcription factor proteins that recognize specific histone-modifying enzymes and recruit them to their sites of function. For example, the transcription factor SUF4 (SUPPRESSOR OF FRIGIDA4) likely recruits the histone methyltransferase EFS (EARLY FLOWERING IN SHORT DAYS) and the PAF1-like complex to the floral inhibitor *FLC* (FLOWERING LOCUS C) promoter (28, 29). The transcription factor ALF (ABI3-like factor) is thought to recruit a histone acetyltransferase (HAT) activity to the *phaseolin* promoter in the bean (*Phaseolus vulgaris*) (30). Transcription factors BPC1 (BASIC PENTACYSTEINE 1), BPC6, and AZF1 (ARABIDOPSIS ZINC FINGER 1) interact with the components of Polycomb repressive complexes PRC2 and PRC1 and recruit them to the GAGA and telobox motifs to promote histone 3 (H3) trimethylation and H2A monoubiquitylation (31, 32). Transcription factors VAL1 (VP1/ABI3-like 1) and VAL2 interact with the PRC1 component LHP1 and recruit it to the BY motifs (33) whereas NAC (NAM, ATAF, CUC) transcription factors NAC050 and NAC052 interact with and recruit the histone demethylase JMJ14 to its target genes (34). This relatively short list of plant histone-modifying enzymes that are recognized and recruited by transcription factors does not include histone deubiquitinases, one of the main histone-modifiers the mechanism of recruitment of which is poorly understood.

Here, we began filling this gap in our knowledge by detecting the specific interaction between the OTLD1 histone deubiquitinase and the LSH10 protein of Arabidopsis. LSH10 belongs to the ALOG family of proteins, such as DUF640 (domain of unknown function 640), known to act as key developmental regulators in land plants. The presence of DNA binding motifs, transcriptional regulation activity, nuclear localization, and homodimerization ability (21, 35-37) suggest that the ALOG proteins may function as specific transcription factors. Our present understanding of the biological functions of different members of the ALOG protein family is very limited. Several *ALOG* or *LSH* (*LIGHT-SENSITIVE SHORT HYPOCOTYLS)* genes have been characterized in Arabidopsis. The initially identified Arabidopsis *LSH* gene is *LSH1*, the protein product of which is involved in the light-dependent regulation of the hypocotyl development (38). Subsequently, LSH3 and LSH4 were shown to suppress the differentiation of organs, such as cotyledons, leaves, and flowers (36, 39, 40), LSH8 was reported to have a role in the regulation of the ABA signaling pathway (41), and LSH9 was shown to regulate hypocotyl elongation by interacting with the temperature sensor ELF3 (42, 43). The function of LSH10, however, remained unknown, and our observations here suggest that it can function as a transcriptional repressor that interacts with the co-repressor histone deubiquitinase OTLD1 and recruit it to promoters of several of its target genes.

The LSH10-OTLD1 interaction was initially detected in yeast and then confirmed by two independent approaches, BiFC and FRET, within living plant cells. In addition to demonstrating the interaction *in planta*, BiFC established the subcellular localization of the LSH10-OTLD1 complexes in the nucleus whereas FRET indicated that, in this complex, the interacting proteins are positioned <10 nm from each other. The likely function of LSH10 in these complexes was investigated using LSH10 loss-of-function and gain-of-function plant lines, both of which indicated that LSH10 acts as a transcriptional repressor and that its target genes include four of the five genes known to be repressed by OTLD1 (15). Specifically, LSH10 repressed the *OSR2, WUS, ABI5*, and *ARL* genes whereas the transcription of the *GA20OX* gene was not affected by LSH10. Consistent with this activity, LSH10 is associated with the chromatin of the promoter regions of each of the *OSR2, WUS, ABI5*, and *ARL* genes in plant cells. In the LSH10 loss-of-function plants, these promoter regions were enriched for monoubiquitylated H2B, suggesting impairment in the OTLD1 recruitment to the target chromatin. Furthermore, LSH10 is also directly bound to the DNA sequences of these promoter regions. The DNA binding activity of LSH10 detected experimentally is supported by the alignment of the secondary structure of LSH10 predicted by AlphaFold (44, 45) (Fig. S2A) with the known DNA binding domain of the Cre recombinase (PDB: 1ouqA) (Fig. S2B); this alignment, using the TM-align algorithm (46), suggests that both structures are in a similar fold with the TM-score of 0.59682, with the LSH10 helices 1-4 merged with the Cre recombinase DNA binding domain helices B-E, respectively (Fig. S2C).

In summary, our observations suggest that LSH10 functions as a specific transcription factor that recognizes OTLD1 and recruits it to the target gene promoters, and that OTLD1 may utilize multiple transcription factors for this purpose, depending on the specific target gene. The latter notion of several, and most likely at least partly redundant pathways for transcriptional control of the OTLD1 target genes—previously suggested to underlie the lack of detectable phenotypes of the *OTLD1* loss-of-function mutants (15)—may explain why both *LSH10* loss-of-function mutants alleles also did not produce discernable morphological or developmental phenotypes (not shown). Note that this study did not endeavor to understand the specific phenotypic effects of LSH10; instead, we aimed to uncover the role of LSH10 as a transcription factor that recognizes and recruits OTLD1 to the target gene promoters as a paradigm for the mechanism by which plant co-repressor histone deubiquitinases are recruited to the target chromatin by their cognate DNA binding repressors.

## Methods

### Plants and growth conditions

The *Arabidopsis thaliana* loss-of-function T-DNA insertional mutants of *LSH10* (*lsh10-1*, SALK_006965; *lsh10-2*, SK14678) were obtained from ABRC (www.arabidopsis.org/abrc/). For genetic complementation, gain-of-function lines of the *lsh10-1* mutant, the Arabidopsis *LSH10* cDNA was amplified using primers listed in Table S1, cloned into pDONR207 (#12213013, Invitrogen) by the BP reaction using the Gateway BP Clonase II (#11789100, Invitrogen), and transferred into the destination vector pMDC32 (47) by the LR reaction using the Gateway LR Clonase II (#11791020, Invitrogen). The recombinant plasmids were subsequently introduced into *Agrobacterium tumefaciens* strain GV3101, and transformed into the *lsh10-1* mutant plants by the floral dip method (48). The transgenic plants were selected on MS medium supplemented with hygromycin (30 mg/l) and timentin (100 mg/l) and confirmed by PCR and RT-qPCR using primers listed in Table S1.

*Nicotiana benthamiana* plants, wild-type *Arabidopsis thaliana* (ecotype Col-0) plants, and the mutant lines were grown on soil in an environment-controlled chamber at 22°C under long-day conditions (16-h light/8-h dark cycle at 140 μE sec^-1^m^-2^ light intensity) as described previously (14, 15).

### Bimolecular fluorescence complementation (BiFC) assay and subcellular localization assay

Arabidopsis *OTLD1, LSH10, LSH4* cDNAs, and the *Tobacco mosaic virus* methyltransferase domain (MT) coding sequence were amplified using primers listed in Table S1, cloned into pDONR207, and transferred into the destination vectors pGTQL1221YC (#61705, Addgene) and pGTQL1211YN (#61704, Addgene). For BiFC, different combinations of the resulting constructs encoding the nEYFP- and cEYFP-tagged proteins were transiently co-expressed with free CFP in the leaves of 4-8-week-old *N. benthamiana* plants by agroinfiltration. Free CFP was expressed from pPZP-RCS2A-DEST-ECFP-C1 (49). For subcellular localization, LSH10 was fused with CFP by transferring its coding sequence by Gateway cloning from pDONR207 into pPZP-RCS2A-DEST-ECFP-N1 (49). The fluorescence signal was recorded 2 days after infiltration using a confocal laser scanning microscope (LSM 900, Zeiss, Germany) with a 40X oil immersion objective and CFP and YFP filters.

### Fluorescence resonance energy transfer (FRET) assay

The coding sequences of *LSH10* and *OTLD1* were transferred by Gateway cloning from pDONR207 into pPZP-RCS2A-DEST-EGFP-N1 (49) and pPZP-RCS2A-DEST-ERFP-N1 (49) to generate the donor vector *p35S::LSH10-GFP* and the acceptor vector *p35S::OTLD1-RFP*, respectively. For positive control, ERFP was amplified from pPZP-RCS2A-DEST-ERFP-N1 using primers listed in Table S1, cloned into pDONR207, and transferred into pPZP-RCS2A-DEST-EGFP-N1 to generate the RFP-GFP fusion construct. These vectors were transiently expressed in the leaves of 4-8-week-old *N. benthamiana* plants via agroinfiltration. The FRET signal was detected and recorded by confocal microscopy using a 40X oil immersion objective.

#### Sensitized emission

A set of three confocal images of the same field of view was taken using the following channel settings: the GFP channel for excitation and emission of the donor chromophore (excitation lasers: 405 nm, emission filter: 400-597 nm), the RFP channel for excitation and emission of the acceptor chromophore (excitation lasers: 561 nm, emission filter: 400-597 nm), and the FRET channel for excitation of the donor and emission of the acceptor chromophores (excitation lasers: 405 nm, emission filter: 597-617 nm). The ImageJ plug-in PixFRET software was used to generate corrected images of SE-FRET efficiency after subtraction of spectral bleed-through.

#### Acceptor photobleaching

In this approach, the emission of the donor fluorophore is compared before and after photobleaching of the acceptor. Photobleaching of the acceptor leads to an increase in the donor fluorescence if any interactions leading to energy transfer occur because it is no longer quenched by the acceptor. Acceptor photobleaching was performed with 100% intensity of lasers at 561 nm, duration of 30 s, 150 interactions for area bleach, and started after 5 images. Images in the acceptor channel (RFP) and donor channel (GFP) were captured simultaneously before and after photobleaching. %AB-FRET was calculated as the percent increase in GFP emission after RFP photobleaching using the following formula: %AB-FRET = [(GFPpost -GFPpre)/GFPpre]x100, where GFPpost is GFP emission after RFP photobleaching, and GFPpre is GFP emission before RFP photobleaching. The %AB-FRET was determined in regions of interest drawn around the entire area of the cell nucleus.

### DNA, RNA extraction, and reverse transcription-quantitative PCR (RT-qPCR)

Plant genomic DNA was extracted using the DNeasy-like Plant DNA Extraction Protocol. RNA from plants was extracted using TRIzol (#15596026, Invitrogen) according to the manufacturer’s instructions. The RNA was used as a template for cDNA synthesis *via* the RevertAid Reverse Transcription Kit (#K1691, Thermofisher) and Hexa-random primers. RT-qPCR was performed using Power SYBR Green PCR master mix (Catalog **#**: 4367659) (Applied Biosystems by Thermo Fisher Scientific, USA) and a QuantStudio™ 3 Real-Time PCR System (Applied Biosystems by Thermo Fisher Scientific). The thermocycler program consisted of pre-denaturation at 95°C for 10 min followed by 40 cycles at 95°C for 15 s and 60°C for 1 min. Each sample was analyzed in 8 biological replicates and 3 technical repeats for each, and the data were calculated by the 2^-ΔΔCt^ formula (50). *A. thaliana Actin, Sand*, and *EF1a* were used as reference genes. Specific primers used in these experiments are detailed in Table S1.

### qChIP and statistical analysis

Approximately 0.8-1g of leaves of 4-week-old *lsh10-1/LSH10-His6* and wild-type plants were harvested and cross-linked with 1% formaldehyde (v/v) for quantitative chromatin immunoprecipitation (qChIP) assays. The qChIP experiments were performed using an EpiQuikTM Plant ChIP Kit (EpiGentek, P-2014-48). According to the manufacturer’s instructions, the chromatin from the plant cells was extracted and sheared; the length of sheared DNA fragments was 200-1000 bp. 5 μl of the crude chromatin extracts were saved for use as an input control, and 100 μl were added into the microwell immobilized with the antibodies (non-specific rabbit IgG Isotype Control (Invitrogen, WB317638) as the negative control, specific rabbit His-tag antibody (GenScript, A00174-40) or anti-monoubiquityl-histone H2B (Lys-120) (5546S, Cell Signaling Technology, Inc.). DNA was released from the antibody-captured protein-DNA complex, reversed, purified through the specifically designed F-Spin Column, and then amplified by qPCR as described above. Input Ct values were adjusted for the dilution factor and ΔCt was calculated by normalizing Ct values to the adjusted input based on the equation ΔCt=Ct(input)-Ct(IP). For anti-His6 qChIP, the % input was calculated as % input=100×2e-ΔCt; fold enrichment was calculated using % input of IP/IgG as [ΔCt(IP)/ΔCt(IgG)] (51), and non-specific background immunosignal control, obtained with the wild-type *Arabidopsis* plants that do not express the His6 epitope, was subtracted from the qPCR data (17). For anti-H2Bub qChIP, the relative fold change was calculated using the 2^-ΔΔCt^ formula and normalized with adjusted input. Statistical significance was determined by paired two-tailed Student’s t-test, with p-values < 0.05, 0.01, or 0.001 corresponding to the statistical probability of >95%, 99%, or 99.9%, respectively, considered statistically significant.

### Electrophoretic mobility shift assays (EMSAs)

The full-length cDNA of *LSH10* was cloned into the GST fusion vector pGEX-5X-1 and propagated in the LEMO21 strain of *Escherichia coli*. The recombinant protein GST-LSH10 was purified using Glutathione Sepharose 4B beads (GE Healthcare, 17-0756-01) according to the manufacturer’s protocol. The probes were designed by two iterations of motif-based sequence analysis of the intergenic regions of the *ARL, WUS, ABI5*, and *OSR2* genes for conserved motifs using the Multiple Expectation Maximizations for Motif Elicitation (MEME) tool (https://meme-suite.org/meme/). The biotin-labeled and unlabeled oligonucleotides corresponding to both strands of the selected sequences (see Fig. 5A and Table S1) were synthesized by Integrated DNA Technologies (Coralville, IA). The oligonucleotides were annealed, and the resulting probes (10 ng) were incubated with the purified protein (1 μg) and dI.dC (1 μg) at room temperature for 20 min in the binding buffer (100 mM Tris, 500 mM KCL, 10 mM DTT, pH 7.5). For competition experiments, a 300-fold molar excess of each unlabeled probe was included in the binding reaction. EMSA was performed using the Light Shift Chemiluminescent EMSA kit (Thermo Scientific) according to the manufacturer’s instructions. The electrophoretic migration of biotin-labeled probes was resolved on 6% native polyacrylamide gels and detected using an enhanced chemiluminescence substrate (Thermo Scientific).

## Supporting information

full supplemental material

## Acknowledgment

The work in the V.C. laboratory was supported by grants from NIH (R35GM144059 and R01GM50224), NSF (MCB1913165 and IOS1758046), and BARD (IS-5276-20) to V.C. and M.L.

## Author Contributions

M.S.V.P. conducted the experiments and analyzed the experimental data. P. T. T. conducted FRET. I.K. initiated the identification of the *LSH10* gene. M.S.V.P. and V.C designed the experiments and, with the help of M.L., wrote, reviewed, and edited the manuscript.

