## Supplementary material for "*Arabidopsis* LSH10 transcription factor interacts with the co-repressor histone deubiquitinase OTLD1 to recruit it to the target promoters": full supplemental material

**Supplementary Information**

**Legends to Supplementary Figures**

**Fig. S1.** Evolutionary analysis of the Arabidopsis LSH protein family. The evolutionary history of Arabidopsis LSH proteins (AtLSH) was inferred using the Maximum Likelihood method and JTT matrix-based model (1). The tree with the highest log likelihood (-2787.03) is shown. The percentage of trees in which the associated taxa clustered together is shown next to the branches. Initial tree(s) for the heuristic search were obtained automatically by applying Neighbor-Join and BioNJ algorithms to a matrix of pairwise distances estimated using the JTT model, and then selecting the topology with a superior log likelihood value. A discrete Gamma distribution was used to model evolutionary rate differences among sites [5 categories (+G, parameter = 0.5059)]. The tree is drawn to scale, with branch lengths measured in the number of substitutions per site. This analysis involved 10 amino acid sequences, identified by the name of the protein and the corresponding AGI locus code; LSH10 is highlighted with a red dot. There was a total of 225 positions in the final dataset. Evolutionary analyses were conducted in MEGA X (2). Scale bar, 0.2 amino acid substitutions per site.

**Fig. S2.** Alignment of the predicted secondary structure of LSH10 and the N-terminal DNA binding domain of the Cre recombinase. (**A**) The ribbon diagram of LSH10 predicted by AlphaFold (AF-Q9S7R3). Helices 1, 2, 3, 4, and zinc ribbon insert conserved in the ALOG domain are colored in cyan, red, green, magenta, and yellow, respectively. (**B**) The ribbon diagram of the Cre recombinase N-terminal DNA binding domain (PDB: 1ouqA) with its helices A-E is colored in cornflower blue. (**C**) Structure alignment between LSH10 and the Cre recombinase N-terminal DNA binding domain. Helices 1, 2, 3, and 4 of LSH10 align with helices B, C, D, and E of the Cre recombinase, respectively.

**References to Supplementary Figure Legends**

1. Jones, D.T., Taylor, W.R., and Thornton, J.M. (1992) The rapid generation of mutation data matrices from protein sequences. *Comput. Appl. Biosci.* **8**, 275-282.

2. Kumar, S., Stecher, G., Li, M., Knyaz, C., and Tamura, K. (2018) MEGA X: molecular evolutionary genetics analysis across computing platforms. *Mol. Biol. Evol.* **35**, 1547-1549.

**Supplementary Tables**

**Table S1**. Primers used for DNA amplification, molecular cloning, and detection.

| No. | Primer name | Sequence (5′ to 3′) | Purposes |
|---|---|---|---|
| 1 | attB1-AtOTLD1 Fw | ggggacaagtttgtacaaaaaagcaggctcaatgactcggattttggttcaaag | Cloning of AtOTLD1 to entry vector pDNOR207 for BiFC and FRET |
| 2 | attB2-AtOTLD1 Rv | ggggaccactttgtacaagaaagctgggtgttccgtggctttgcctttgcgtc | Cloning of AtOTLD1 to entry vector pDNOR207 for BiFC and FRET |
| 3 | attB1-AtLSH10 Fw | ggggacaagtttgtacaaaaaagcaggctcaatgtcctctccaagagaaagagg | Cloning of AtLSH10 to entry vector pDNOR207 for BiFC, FRET, and genetic complementation |
| 4 | attB2-AtLSH10 Rv | ggggaccactttgtacaagaaagctgggtgatgtcaacagagactaaagaaac | Cloning of AtLSH10 to entry vector pDNOR207 for BiFC and FRET |
| 5 | attB2-His6-linker-AtLSH10 Rv | ggggaccactttgtacaagaaagctgggtgtcagtgatggtgatggtgatgcccaccaccagagaagctgaaggaagaggagg | Cloning of AtLSH10 to entry vector pDNOR207 for genetic complementation |
| 6 | attB1-AtLSH4 Fw | ggggacaagtttgtacaaaaaagcaggctcaatggatcatatcatcggctttatg | Cloning of AtLSH4 to entry vector pDNOR207 for BiFC |
| 7 | attB2-AtLSH4 Rv | ggggaccactttgtacaagaaagctgggtgattagggctacttgaaatcgcc | Cloning of AtLSH4 to entry vector pDNOR207 for BiFC |
| 8 | attB1-TMV MT Fw | ggggacaagtttgtacaaaaaagcaggctc | Cloning of TMV MT to entry vector pDNOR207 for BiFC |
| 9 | attB2-TMV MT Rv | ggggaccactttgtacaagaaagctgggtg | Cloning of TMV MT to entry vector pDNOR207 for BiFC |
| 10 | AttB1 mRFP Fw | ggggacaagtttgtacaaaaaagcaggctcaatggcctcctccgaggacgt | Cloning of mRFP to entry vector pDNOR207 for FRET |
| 11 | AttB2 mRFP Rv | ggggaccactttgtacaagaaagctgggtgttggagatctgcggccgcgg | Cloning of mRFP to entry vector pDNOR207 for FRET |
| 12 | AttL1 | tcgcgttaacgctagcatggatctc | Confirmative PCR and sequencing in entry clones |

| | | | |
|---|---|---|---|
| 13 | AttL2 | gtaacatcagagattttgagacac | Confirmative PCR and sequencing in entry clones |
| 14 | AttB1 Fw | ggggacaagtttgtac aaaaaagcaggct | Confirmative PCR and sequencing in destination clones |
| 15 | AttB2 Rv | ggggaccactttgta caagaaagctgggt | Confirmative PCR and sequencing in destination clones |
| 16 | 35S Promoter Fw | ctatccttcgcaagacccttc | Confirmative PCR in destination binary vector |
| 17 | GFP 86 Rv | ctgaacttgtggccgtttacgt | Confirmative PCR in BiFC vector with nYFP |
| 18 | GFP 604 Rv | ggtagtggttgtcgggcag | Confirmative PCR in BiFC vector with cYFP |
| 19 | LSH10-1 Fw | ccgttacgagtcgcagaaga | genotyping of Arabidopsis *lsh10-1* mutant line, gene specific |
| 20 | LSH10-1 Rv | accggtcccttaattccgaac | genotyping of Arabidopsis *lsh10-1* mutant line, gene specific |
| 21 | SALK_LBb1.3 | attttgccgatttcggaac | genotyping of Arabidopsis *lsh10-1* mutant line, Ti-DNA specific |
| 22 | LSH10-2 Fw | tcgtggtggagaactcaaact | genotyping of Arabidopsis *lsh10-2* mutant line, gene specific |
| 23 | LSH10-2 Rv | ccccaagcttgtctgagagg | genotyping of Arabidopsis *lsh10-2* mutant line, gene specific |
| 24 | SK LB R | tctaagccccccatttggacg | genotyping of Arabidopsis *lsh10-2* mutant line, Ti-DNA specific |
| 25 | AtActin2 exon-span 1093 Fw | gtggtcgtacaaccggtatt | Reference for qRT-PCRs of transcripts in Arabidopsis, span exon-exon boundary to avoid genomic DNA contamination |

| | | | |
|---|---|---|---|
| 26 | AtActin2 1288 Rv | cacgtccagcaaggtcaaga | Reference for qRT-PCRs of transcripts in Arabidopsis |
| 27 | AtSand Fw | gctgatgatggaagcgagga | Reference for qRT-PCRs of transcripts in Arabidopsis, span exon-exon boundary to avoid genomic DNA contamination |
| 28 | AtSand exon-span Rv | tgacgtagaagcatcatcctcat | Reference for qRT-PCRs of transcripts in Arabidopsis |
| 29 | AtEF1a exon-span Fw | atgatttgctgttgtaacaagatgga | Reference for qRT-PCRs of transcripts in Arabidopsis, span exon-exon boundary to avoid genomic DNA contamination |
| 30 | AtEF1a Rv | ggcttgtctgatggcctctt | Reference for qRT-PCRs of transcripts in Arabidopsis |
| 31 | AtLSH10 exon-span Fw | gagatgggaggtgggagaga | qRT-PCRs of AtLSH10 transcripts, span exon-exon boundary to avoid genomic DNA contamination |
| 32 | AtLSH10 Rv | ctgatcctgccaagttcaca | qRT-PCRs of AtLSH10 transcripts |
| 33 | AtABI5 exon-span Fw | ccgcgagtctgctgctagat | qRT-PCRs of AtABI5 transcripts, span exon-exon boundary to avoid genomic DNA contamination |
| 34 | AtABI5 Rv | ttcctctccaactccgccaat | qRT-PCRs of AtABI5transcripts |
| 35 | AtARL exon-span Fw | gctctaacggtttaaagctttctt | qRT-PCRs of AtARL transcripts, span exon-exon boundary to avoid genomic DNA contamination |
| 36 | AtARL Rv | tcttggcgtttccagaccta | qRT-PCRs of AtARL transcripts |
| 37 | AtWUS exon-span Fw | tcccagcttcaataacgggaat | qRT-PCRs of AtWUS transcripts, span exon-exon boundary to avoid genomic DNA contamination |

| | | | |
|---|---|---|---|
| 38 | AtWUS Rv | tcctccacctacgttgttgtaat | qRT-PCRs of AtWUStranscripts |
| 39 | AtOSR2 Fw | tgatggtgctattggcggttct | qRT-PCRs of AtOSR2 transcripts |
| 40 | AtOSR2 Rv | agcattagcatcagcaccaccg | qRT-PCRs of AtLSH10 transcripts |
| 41 | a- AtOSR2 Fw | taatataaat tgcgttttgttttc cagatatcaa | Biotin labeled and unlabeled oligos for EMSA, probe a |
| 42 | a- AtOSR2 Rv | ttgatatctggaaaacaaaacgcaatttatatta | Biotin labeled and unlabeled oligos for EMSA, probe a |
| 43 | b- AtOSR2 Fw | ctgtaacgtctttctcttcacacacataaaaaag | Biotin labeled and unlabeled oligos for EMSA, probe b |
| 44 | b- AtOSR2 Rv | cttttttatg tgtgtgagagagaaa gacgttacag | Biotin labeled and unlabeled oligos for EMSA, probe b |
| 45 | c- AtWUS Fw | ttttagaatt tttgttttgttttc tgtgtgtatg | Biotin labeled and unlabeled oligos for EMSA, probe c |
| 46 | c- AtWUS Rv | catacacacagaaaacaaaacaaaaattctaaaa | Biotin labeled and unlabeled oligos for EMSA, probe c |
| 47 | d- AtWUS Fw | tatatatatataatcttctctctcacacaaaacctaaaa | Biotin labeled and unlabeled oligos for EMSA, probe d |
| 48 | d- AtWUS Rv | ttttaggttttgtgtgagagagaagattatatatatata | Biotin labeled and unlabeled oligos for EMSA, probe d |
| 49 | e- AtWUS Fw | atttaccgttaactttgtgaacaaaagtcgaatcaaac acac | Biotin labeled and unlabeled oligos for EMSA, probe e |
| 50 | e- AtWUS Rv | gtgtgtttgattcgacttttgttcacaaagttaacggta aat | Biotin labeled and unlabeled oligos for EMSA, probe e |
| 51 | f- AtABI5 fw | gcttttaaactatgtgaaggaggagaacc tccataacaa | Biotin labeled and unlabeled oligos for EMSA, probe f |
| 52 | f- AtABI5 Rv | ttgttatggaggttctcctccttcacatagtttaaaagc | Biotin labeled and unlabeled oligos for EMSA, probe f |
| 53 | g-AtABI5 Fw | acaagaagcggattctctcagttttccggcgg cggaggaaca | Biotin labeled and unlabeled oligos for EMSA, probe g |
| 54 | g-AtABI5 Rv | tgttcctccgccgccggaaaactgagagaatccgct tcttgt | Biotin labeled and unlabeled oligos for EMSA, probe g |

| | | | |
|---|---|---|---|
| 55 | h-AtARL Fw | tcagagaaagtgtctaggggagaagaaccactgtg attg | Biotin labeled and unlabeled oligos for EMSA, probe h |
| 56 | h-AtARL Rv | caatcacagtggttcttctcccctagacactttctctga | Biotin labeled and unlabeled oligos for EMSA, probe h |
| 57 | j-AtARL Fw | agcaacttgcctctcacacttgtc | Biotin labeled and unlabeled oligos for EMSA, probe j |
| 58 | j-AtARL Rv | gacaagtgtgagaggcaagttgct | Biotin labeled and unlabeled oligos for EMSA, probe j |
| 59 | k-AtOSR2 Fw | agtttaagccgacacgttaatacat | Biotin labeled and unlabeled oligos for EMSA, probe k |
| 60 | k-AtOSR2 Rv | atgtattaacgtgtcggcttaaact | Biotin labeled and unlabeled oligos for EMSA, probe k |
| 61 | l-AtWUS Fw | tagagaggaa caagagaaagagaag tttgaagtta | Biotin labeled and unlabeled oligos for EMSA, probe l |
| 62 | l-AtWUS Rv | tagagaggaacaagagaaagagaagtttgaagtta | Biotin labeled and unlabeled oligos for EMSA, probe l |
| 63 | AtOSR2 pro Fw | ccagcaagttgttttcttgctaac | qChip-PCRs |
| 64 | AtOSR2 pro Rv | tggaagaaccgccaatagca | qChip-PCRs |
| 65 | AtWUS pro Fw | ccagcaagttgttttcttgctaac | qChip-PCRs |
| 66 | AtWUS pro Rv | tgtgtttgattcgacttttgttcac | qChip-PCRs |
| 67 | AtABI5 pro Fw | accgcctcctacccatttatc | qChip-PCRs |
| 68 | AtABI5 pro Rv | aaaaccggtggctttgtgttc | qChip-PCRs |
| 69 | AtARL pro Fw | tcacacttgtcaccccccaaa | qChip-PCRs |
| 70 | AtARL pro Rv | tggggttaagactgtggaatca | qChip-PCRs |

**Table S2.** Raw data for quantification of AB-FRET (for Fig. 2C).

| | % AB-FRET | | | |
|---|---|---|---|---|
| **Cell Images** | **RFP-GFP FUSION** | **LSH10-GFP + OTLD1-RFP** | **LSH4-GFP + OTLD1-RFP** | **LSH10-GFP + FREE-RFP** |
| 1 | 24.62 | 15.84 | 1.76 | -0.17 |
| 2 | 31.91 | 15.87 | -0.21 | -0.55 |
| 3 | 27.34 | 18.27 | 0.32 | -5.83 |
| 4 | 38.26 | 15.68 | -4.11 | -0.64 |
| 5 | 30.86 | 15.85 | -5.29 | 1.78 |
| 6 | 32.34 | 13.58 | -2.51 | -1.39 |
| 7 | 35.49 | 11.50 | -0.12 | -2.64 |
| 8 | 30.03 | 7.97 | -1.15 | -2.38 |
| 9 | 24.86 | 11.31 | 0.22 | -0.13 |
| 10 | 30.49 | 4.40 | -0.28 | 2.16 |
| 11 | 26.38 | 9.73 | -0.80 | 0.49 |
| 12 | 30.08 | 12.39 | -1.38 | 0.04 |
| 13 | 21.78 | 9.31 | 0.00 | -0.44 |
| **Mean** | **29.57** | **12.44** | **-1.04** | **-0.75** |

**Table S3.** Raw data for quantification of the increase in expression of the target genes in the *lsh10-1,* and *lsh10-2* plants (for Fig. 4B).

| **Mutants** | **Relative fold change** | | | | | |
|---|---|---|---|---|---|---|
| **Samples** | **LSH10** | **OSR2** | **WUS** | **ABI5** | **ARL** | **EF1a** |
| WT1 | 1.39 | 0.28 | 0.18 | 0.36 | 0.49 | 0.90 |
| WT2 | 1.50 | 0.59 | 0.67 | 0.87 | 0.88 | 0.85 |
| WT3 | 0.66 | 0.86 | 0.87 | 1.05 | 0.56 | 1.02 |
| WT4 | 1.16 | 1.76 | 1.97 | 0.83 | 1.26 | 1.10 |
| WT5 | 1.48 | 2.13 | 2.22 | 1.07 | 1.29 | 1.16 |
| WT6 | 0.94 | 0.74 | 0.63 | 1.25 | 0.87 | 1.21 |
| WT7 | 0.50 | 1.06 | 1.84 | 2.08 | 2.22 | 1.05 |
| WT8 | 0.89 | 0.90 | 1.34 | 1.64 | 1.25 | 0.79 |
| **Mean** | **1.07** | **1.19** | **1.28** | **1.11** | **1.10** | **1.01** |
| *lsh10-1-1* | 0.03 | 19.52 | 20.65 | 15.08 | 89.98 | 1.02 |
| *lsh10-1-2* | 0.06 | 14.25 | 32.27 | 19.10 | 31.69 | 0.86 |
| *lsh10-1-3* | 0.06 | 39.52 | 21.97 | 22.50 | 27.36 | 0.97 |
| *lsh10-1-4* | 0.03 | 18.02 | 30.51 | 17.04 | 58.72 | 1.23 |
| *lsh10-1-5* | 0.03 | 16.90 | 16.42 | 5.53 | 34.73 | 1.40 |
| *lsh10-1-6* | 0.04 | 22.89 | 25.78 | 26.34 | 28.37 | 1.03 |
| *lsh10-1-7* | 0.03 | 22.89 | 51.03 | 18.84 | 65.83 | 1.05 |
| *lsh10-1-8* | 0.02 | 44.56 | 17.34 | 38.94 | 49.13 | 1.00 |
| **Mean** | **0.04** | **24.82** | **26.99** | **20.42** | **48.23** | **1.07** |
| *lsh10-2-1* | 0.02 | 76.58 | 25.25 | 5.46 | 63.08 | 0.72 |
| *lsh10-2-2* | 0.01 | 17.46 | 24.20 | 11.08 | 87.23 | 1.11 |

| | | | | | | |
|---|---|---|---|---|---|---|
| *lsh10-2-3* | 0.02 | 35.08 | 16.71 | 9.77 | 129.33 | 0.86 |
| *lsh10-2-4* | 0.01 | 9.39 | 20.51 | 5.62 | 105.41 | 0.86 |
| *lsh10-2-5* | 0.01 | 25.85 | 42.61 | 6.01 | 25.41 | 1.47 |
| *lsh10-2-6* | 0.01 | 11.66 | 9.95 | 10.26 | 30.57 | 1.28 |
| *lsh10-2-7* | 0.02 | 15.25 | 40.48 | 11.59 | 10.43 | 1.29 |
| *lsh10-2-8* | 0.03 | 12.20 | 12.70 | 7.83 | 78.31 | 0.99 |
| **Mean** | **0.01** | **25.43** | **24.05** | **8.45** | **66.22** | **1.07** |

**Table S4.** Raw data for quantification of the transcriptional repression of the target genes in the *lsh10-1/LSH10-His6* plants (for Fig. 4C).

| **Transgenic plants** | **Relative fold change** | | | | | |
|---|---|---|---|---|---|---|
| **Samples** | **LSH10** | **OSR2** | **WUS** | **ABI5** | **ARL** | **EF1a** |
| WT1 | 1.21 | 1.47 | 1.29 | 0.95 | 0.80 | 1.09 |
| WT2 | 0.90 | 0.96 | 1.10 | 0.77 | 1.21 | 1.24 |
| WT3 | 1.13 | 1.25 | 1.65 | 0.79 | 0.72 | 1.13 |
| WT4 | 1.19 | 2.30 | 0.39 | 2.30 | 0.39 | 1.05 |
| WT5 | 0.78 | 0.82 | 0.66 | 0.93 | 1.02 | 1.01 |
| WT6 | 1.05 | 0.69 | 0.65 | 1.87 | 1.40 | 0.97 |
| WT7 | 0.77 | 2.33 | 0.19 | 3.57 | 0.75 | 0.71 |
| WT8 | 1.09 | 1.59 | 1.20 | 2.26 | 0.52 | 0.90 |
| **Mean** | **1.01** | **1.43** | **0.89** | **1.68** | **0.85** | **1.01** |
| *lsh10-1/LSH10-His6-1* | 1.53 | 1.30 | 0.54 | 0.81 | 0.80 | 1.14 |
| *lsh10-1/LSH10-His6-2* | 1.36 | 1.24 | 0.69 | 0.87 | 1.02 | 1.19 |
| *lsh10-1/LSH10-His6-3* | 1.27 | 0.65 | 0.53 | 0.53 | 1.09 | 1.22 |
| *lsh10-1/LSH10-His6-4* | 1.30 | 1.16 | 0.32 | 1.13 | 0.77 | 1.22 |
| *lsh10-1/LSH10-His6-5* | 1.35 | 0.44 | 0.24 | 0.73 | 0.37 | 0.95 |
| *lsh10-1/LSH10-His6-6* | 1.14 | 0.69 | 0.12 | 0.80 | 0.84 | 0.99 |
| *lsh10-1/LSH10-His6-7* | 1.42 | 0.44 | 0.09 | 0.95 | 0.96 | 1.07 |
| lsh10-1/LSH10-His6-8 | 1.00 | 0.66 | 0.42 | 0.88 | 0.34 | 1.18 |
| **Mean** | **1.30** | **0.82** | **0.37** | **0.84** | **0.77** | **1.12** |

**Table S5.** Raw data for quantification of the qChIP analysis of the association of LSH10-His6 with the chromatin of the target genes (for Fig. 6A).

| **Transgenic plants** | **Fold enrichment** | | | |
|---|---|---|---|---|
| **Samples** | **OSR2** | **WUS** | **ABI5** | **ARL** |
| WT1 | 0.38 | 1.01 | 2.03 | 0.86 |
| WT2 | 0.60 | 0.31 | 0.71 | 0.27 |
| WT3 | 0.82 | 0.75 | 0.16 | 0.17 |
| WT4 | 0.31 | 0.52 | 0.21 | 0.18 |
| WT5 | 0.81 | 0.65 | 0.19 | 0.19 |
| Mean | 0.58 | 0.65 | 0.66 | 0.33 |
| *lsh10-1/LSH10-His6-1* | 1.70 | 1.07 | 1.40 | 1.76 |
| *lsh10-1/LSH10-His6-2* | 3.17 | 0.75 | 1.00 | 1.30 |
| *lsh10-1/LSH10-His6-3* | 2.19 | 0.71 | 1.53 | 0.65 |
| *lsh10-1/LSH10-His6-4* | 0.84 | 1.36 | 3.74 | 1.17 |
| *lsh10-1/LSH10-His6-5* | 1.16 | 1.03 | 0.80 | 0.80 |
| *lsh10-1/LSH10-His6-6* | 0.56 | 2.02 | 2.85 | 2.24 |
| *lsh10-1/LSH10-His6-7* | 0.93 | 2.53 | 3.96 | 4.31 |
| **Mean** | **1.51** | **1.35** | **2.18** | **1.75** |
| **Transgenic plants** | **LSH10-His6 (A.U.)** | | | |
| **Samples** | **OSR2** | **WUS** | **ABI5** | **ARL** |
| *lsh10-1/LSH10-His6-1* | 1.12 | 0.42 | 0.74 | 1.43 |
| *lsh10-1/LSH10-His6-2* | 2.58 | 0.10 | 0.34 | 0.97 |
| *lsh10-1/LSH10-His6-3* | 1.60 | 0.06 | 0.86 | 0.32 |

| | | | | |
|---|---|---|---|---|
| *lsh10-1/LSH10-His6-4* | 0.26 | 0.71 | 3.08 | 0.83 |
| *lsh10-1/LSH10-His6-5* | 0.58 | 0.38 | 0.14 | 0.47 |
| *lsh10-1/LSH10-His6-6* | -0.02 | 1.37 | 2.18 | 1.90 |
| *lsh10-1/LSH10-His6-7* | 0.34 | 1.88 | 3.30 | 3.97 |
| **Mean** | **0.92** | **0.70** | **1.52** | **1.41** |

**Table S6.** Raw data for quantification of the qChIP analysis of the increase in H2B monoubiquitylation of the target chromatin (for Fig. 6B).

| **Mutants** | **Relative fold enrichment (H2B-Ub)** | | | |
|---|---|---|---|---|
| **Samples** | **OSR2** | **WUS** | **ABI5** | **ARL** |
| WT1 | 0.71 | 1.18 | 0.77 | 0.68 |
| WT2 | 0.78 | 1.09 | 0.66 | 0.85 |
| WT3 | 1.36 | 0.83 | 1.40 | 1.33 |
| WT4 | 1.34 | 0.92 | 1.41 | 1.29 |
| WT5 | 1.38 | 1.14 | 2.48 | 2.48 |
| WT6 | 0.82 | 0.81 | 1.53 | 1.75 |
| WT7 | 0.88 | 1.12 | 0.26 | 0.23 |
| **Mean** | **1.04** | **1.01** | **1.22** | **1.23** |
| *lsh10-1-1* | 3.65 | 1.70 | 8.15 | 3.03 |
| *lsh10-1-2* | 3.49 | 6.07 | 2.20 | 2.03 |
| *lsh10-1-3* | 3.44 | 1.22 | 3.19 | 3.80 |
| *lsh10-1-4* | 2.03 | 2.21 | 0.62 | 1.77 |
| *lsh10-1-5* | 5.61 | 1.34 | 5.12 | 3.84 |
| *lsh10-1-6* | 7.22 | 3.15 | 8.24 | 3.62 |
| **Mean** | **4.24** | **2.62** | **4.59** | **3.02** |
| *lsh10-2-1* | 2.59 | 1.56 | 1.32 | 2.33 |
| *lsh10-2-2* | 1.87 | 2.81 | 7.88 | 3.14 |
| *lsh10-2-3* | 2.11 | 1.52 | 7.61 | 2.97 |
| *lsh10-2-4* | 3.28 | 2.11 | 2.24 | 3.33 |

| | | | | |
|---|---|---|---|---|
| *lsh10-2-5* | - | 2.78 | 2.86 | 3.98 |
| *lsh10-2-6* | - | 0.94 | 2.96 | 2.03 |
| *lsh10-2-7* | - | 1.31 | 4.88 | 1.19 |
| **Mean** | **2.46** | **1.86** | **4.25** | **2.71** |

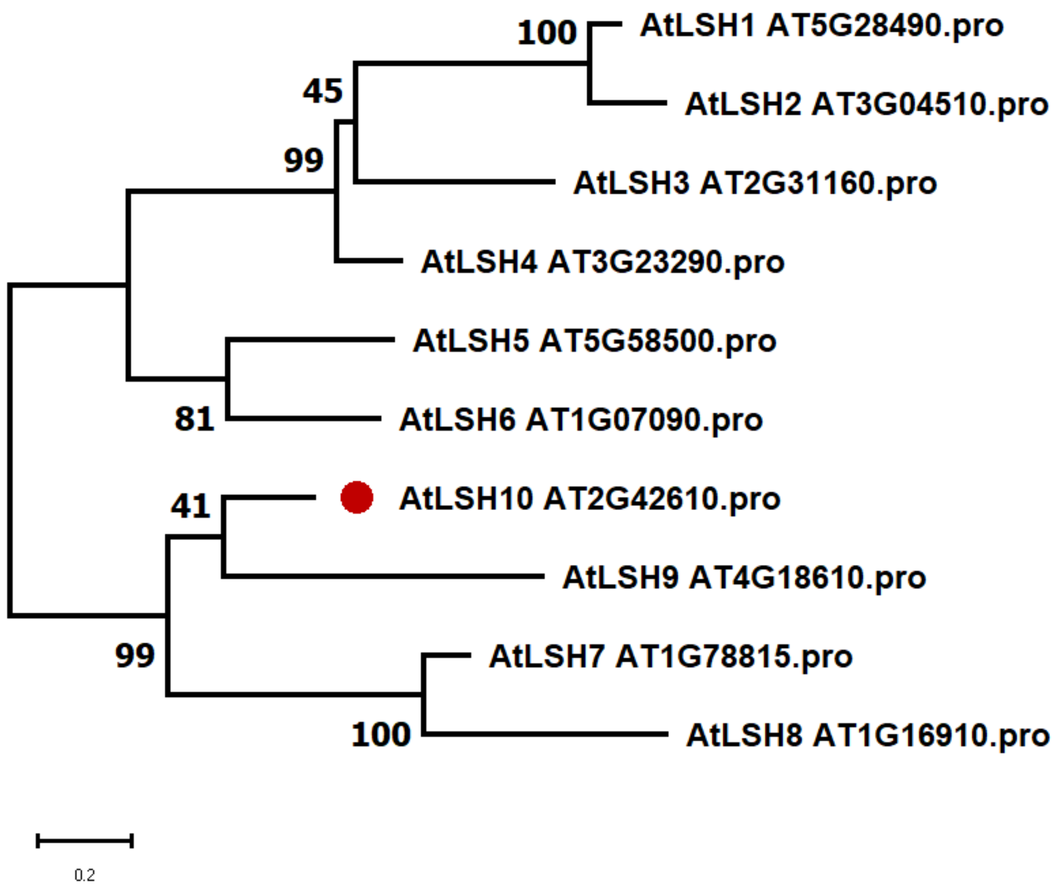

100
AtLSH1 AT5G28490.pro
45
AtLSH2 AT3G04510.pro
99
AtLSH3 AT2G31160.pro
AtLSH4 AT3G23290.pro
AtLSH5 AT5G58500.pro
81
AtLSH6 AT1G07090.pro
41
AtLSH10 AT2G42610.pro
AtLSH9 AT4G18610.pro
99
AtLSH7 AT1G78815.pro
100
AtLSH8 AT1G16910.pro
0.2

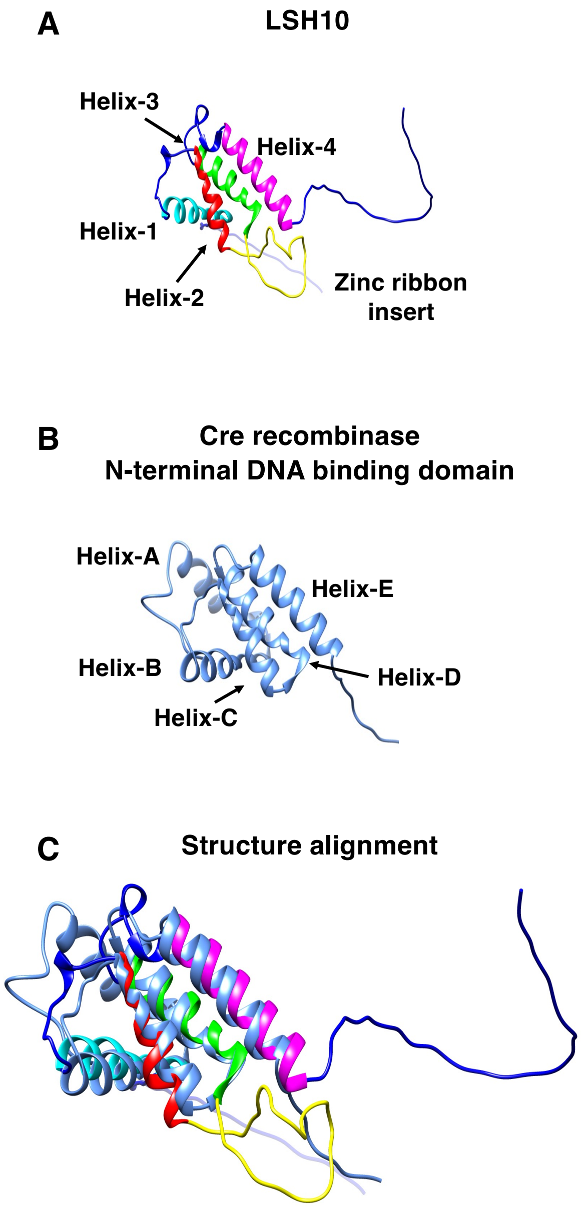
A
LSH10
Helix-3
Helix-4
Helix-1
Helix-2
Zinc ribbon insert
B
Cre recombinase
N-terminal DNA binding domain
Helix-A
Helix-E
Helix-B
Helix-D
Helix-C
C
Structure alignment
